# Comparing management strategies for conserving communities of climate-threatened species with a stochastic metacommunity model

**DOI:** 10.1101/2021.12.01.470788

**Authors:** Gregory A. Backus, Yansong Huang, Marissa L. Baskett

## Abstract

Many species are shifting their ranges to keep pace with climate change, but habitat fragmentation and limited dispersal could impede these range shifts. In the case of climate-vulnerable foundation species such as tropical reef corals and temperate forest trees, such limitations might put entire communities at risk of extinction. Restoring connectivity through corridors, stepping-stones, or enhanced quality of existing patches could prevent the extinction of several species, but dispersal-limited species might not benefit if other species block their dispersal. Alternatively, managers might relocate vulnerable species between habitats through assisted migration, but this is generally a species-by-species approach. To evaluate the relative efficacy of these strategies, we simulated the climate-tracking of species in randomized competitive metacommunities with alternative management interventions. We found that corridors and assisted migration were the most effective strategies at reducing extinction. Assisted migration was especially effective at reducing the extinction likelihood for short-dispersing species, but it often required moving several species repeatedly. Assisted migration was more effective at reducing extinction in environments with higher stochasticity, and corridors were more effective at reducing extinction in environments with lower stochasticity. We discuss the application of these approaches to an array of systems ranging from tropical corals to temperate forests.

## Introduction

The projected rate of climate change threatens many species, especially dispersal-limited species (Urban 2015). Habitat fragmentation intensifies this risk by causing the additional impediment of needing to disperse over poor-quality habitat (Krosby et al. 2010). Moreover, when competing species track climate change at differential speeds, faster dispersing species can block slower dispersing species from tracking climate change (Urban et al. 2012). Such impediments can have ecosystem-wide consequences when dispersal-limited species serve as foundation species, such as in forests (Honnay et al. 2002) and tropical coral reefs (Munday et al. 2009). Though many coral reef species can disperse far in their larval stage, differential dispersal ability and fragmentation could mean that some species are unable to keep pace with climate change (Gaines et al. 2007; Munday et al. 2009). Similarly, competition and the differential effects of climate change on tree species means that poleward species might prevent equatorward species from tracking climate change, especially over fragmented landscapes (Scheller & Mladenoff 2008).

One potential method of conserving dispersal-limited species is through assisted migration, or the relocation of populations outside of the species’ historical range to areas that will be more suitable in response to climate change (McLachlan et al. 2007; Hoegh-Guldberg et al. 2008). Despite a long history of conservation translocations within a species’ historical range (Seddon et al. 2007), relocating a species to a new area with novel species interactions could pose additional challenges. With little precedent and high uncertainty, relocated populations could become invasive or spread diseases and parasites (Mueller & Hellmann 2008; Ricciardi & Simberloff 2009). Even translocations within a species’ range are often unsuccessful without the additional complications of novel competitors, climate change, and fragmentation (Fischer & Lindenmayer 2000; Bubac et al. 2019). To limit relocation failure, decision-making frameworks for assisted migration generally seek to understand the uncertainty around the optimal time and place to move a vulnerable species (McDonald-Madden et al. 2011). However, assisted migration might have limited success when relocating species with narrow climate tolerance into environments with high climate variability over time or low climate variability over space. Additionally, assisted migration is often a single-species approach (Lawler & Olden 2011) that addresses the symptoms of extinction risk instead of the root causes (e.g., habitat fragmentation; Fazey & Fischer 2009; Minteer & Collins 2010). Despite potential risks and uncertainties, assisted migration is already underway for several species at risk of extinction (Liu et al. 2012; Seddon et al. 2015; Wang et al. 2019) and some variations of assisted migration are being tested in coral reefs (van Oppen et al. 2014; Kuffner et al. 2020) and trees (Sáenz-Romero et al. 2021).

Alternatively, habitat restoration in and between fragmented habitats could assist the natural dispersal of species that would otherwise be unable to track climate change (Lawler & Olden 2011). Building habitat corridors (Beier & Noss 1998; Haddad et al. 2015) or stepping-stones reserves (McDowell et al. 1991; Treml et al. 2008) might help increase connectivity and decrease extinction risk from climate change (Robillard et al. 2015), and additional protection of existing reserves might bolster source populations to increase overall persistence (Heller and Zavaleta 2009). Unlike the single-species focus of assisted migration, increasing habitat protection or connectivity is a community-level approach that could directly benefit multiple species that might otherwise be unable to disperse between fragmented patches (Lawler & Olden 2011). However, increasing connectivity and habitat protection do not specifically target species disproportionally affected by climate change, where biological limitations in dispersal ability and negative effects of community interactions could prevent climate tracking (Gilman et al. 2010; Urban et al. 2012). Among the restoration options, those that increase connectivity inherently increase available habitat area, which could be critical for declining populations at risk of extinction from climate change (Hodgson et al. 2009). While increasing connectivity typically has a smaller effect on population outcomes than increasing protection or patch size, or reducing overall habitat loss, in conservation generally (Harrison & Bruna 1999; Fahrig 2001; Falcy & Estades 2007), increasing connectivity might have a greater impact when considering range shift dynamics under climate change (Nuñez et al. 2013). Like assisted migration, the effectiveness of connectivity and restoration-based approaches at conserving species can depend on spatio-temporal variability, as stochasticity in connectivity can reduce species’ persistence (Watson et al. 2012) while heterogeneity in microclimates can increase persistence through climate change (Suggitt et al 2018). As an example of a connectivity-based approach, protecting a marine reserve network focused on connectivity between locations with different levels of temperature stress is one proposed approach to buffer coral reef response to climate change (Mumby et al. 2011; Walsworth et al. 2019). For forest trees, connectivity and restoration would involve creating large-scale networks of land-sharing or land-sparing between disconnected forests (Fischer et al. 2014) or working with local landowners to encourage practices that reduce barriers and promote species persistence (Krosby et al. 2010).

Given the potential trade-offs to each approach, we compare the relative efficacy of these alternative management strategies to support species responses to climate change. To understand how these strategies compare under a variety of conditions in terms of spatio-temporal climate variability, we built a metacommunity model that simulates climate tracking of several randomized species competing in a fragmented environment over a temperature gradient through a cycle of reproduction, dispersal, and competition. Using this model, we compared a variety of management strategies to conserve species’ persistence and diversity: assisted migration, building habitat corridors, creating stepping-stone reserves, and reinforcing areas that currently had high habitat quality.

## Methods

To compare the potential for various conservation strategies to reduce extinction in environments under different spatio-temporal conditions, we modeled metacommunity dynamics of species competing on a one-dimensional linear temperature gradient subjected to climate change. Building on the models by Backus & Baskett (2021) and Urban et al. (2012), all species in this metacommunity compete for the same resources on the same trophic level; we focus on competition as the central interspecific interaction because of its role in range limits (Connell 1972, Sexton et al. 2009) and range shifts. Each species *i* has a discrete population size *n*_*i*_(*x,y,t*) that changes with time *t* and space on both the large *x* and local scale *y*. All populations cycle through reproduction, dispersal, and competition, each with demographic stochasticity. Each species has a unique thermal optimum *ζ*_*i*_ dispersal distance *γ*_*i*_, thermal tolerance breadth *σ*_*i*_, and reproductive strength *ρ*_*i*_. The carrying capacity *K*(*x,y*) varies over space to represent high- and low-quality habitat. After simulating metacommunity dynamics with climate change, we compared extinction rates under each approach. Then we focused on comparing corridors to assisted migration for different levels of environmental stochasticity and local heterogeneity, and finally we analyzed the species characteristics associated with protection by each approach.

### Climate variability and change

We represent local temperature variation across space with the local climate heterogeneity parameter, *H*. Space in this model is a one-dimensional temperature gradient of *L* patches, representing large-scale latitudinal or elevational change (Urban et al. 2012). Each patch *x* ∈ *X* has *W* subpatches, representing small-scale variability in microclimates without an explicit spatial structure. Each local subpatch *y* ∈ *Y* temperature has *T*(*x,y,t*) with a mean patch temperature of 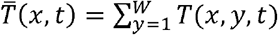 at time *t*.We set the local climate heterogeneity such that each patch has a standard deviation in local temperatures of

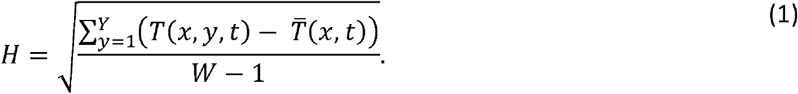

Temperature increases linearly over time with environmental stochasticity, *S*, representing the magnitude of interannual variation in temperature across the environment. At the beginning of each time step, all patches simultaneously increase in temperature by an average value of *τ*, with a stochastic component with autocorrelation *k*, and standard deviation *S* around white noise 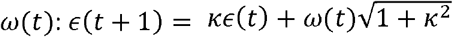, with the square root term to remove the effect of autocorrelation on the variance (Wichmann et al. 2005). Altogether, the temperature in patch *x* changes over time is

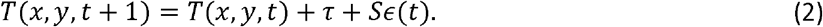

### Metacommunity dynamics

Each simulated species *i* has a population size population size of *n*_*i*_(*x*.*y,t*) individuals in patch *x*, subpatch *y* at discrete time *t*. All individuals reproduce simultaneously at the beginning of each time step with a reproductive output *b*_*i*_(*T*(*x,y,t*)) as a function of time- and location-dependent temperature (Fig. 1a). Temperature-dependence is skew-normal, given skewness constant *λ* with the highest values around the species’ thermal optimum *ζ*_*i*_ and a sharp decrease above *ζ*_*i*_ (Norberg 2004). Then given, thermal tolerance breadth *σ*_*i*_ and fecundity *ρ*_*i*_, the reproductive output is

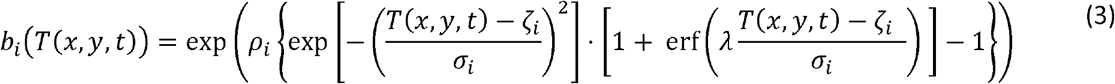

**Figure 1:**
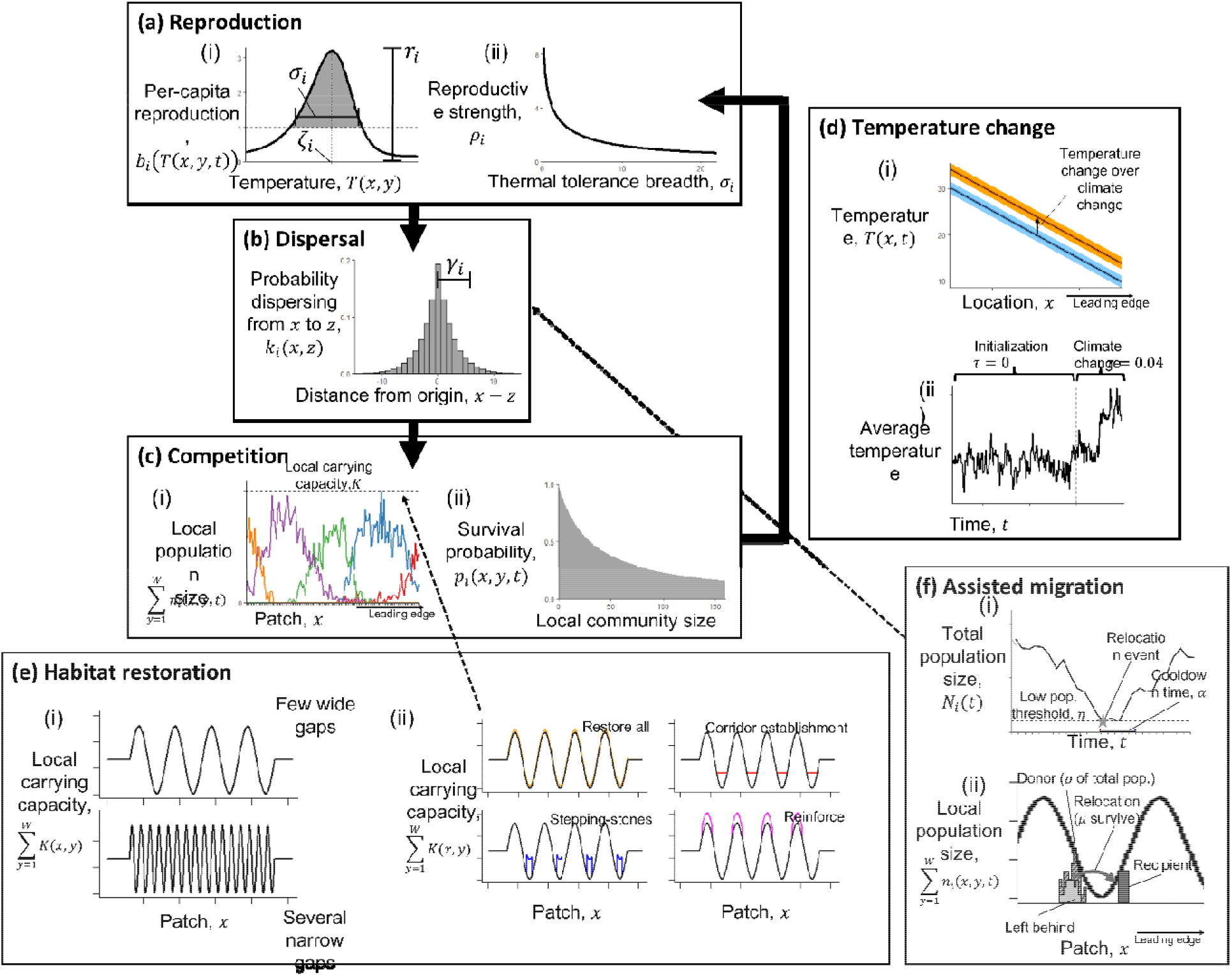
Figure 1: During each time step of the model, all extant species cycle through (a) reproduction, (b) dispersal, and (c) competition before (d) the temperature changes and the next time step continues. (a.i) Per capita reproductive output is skew-normal, dependent on temperature This function is shaped by species’ thermal optimum and thermal tolerance breadth. (a.ii) Reproductive strength scales the total reproductive output so that species with narrow (specialists) have higher reproduction and species with broad (generalists) have lower reproduction. (b) The dispersal kernel is a long-tailed “double geometric” distribution with a mean dispersal distance. (c.i) All species compete over limited space, where each patch has a carrying capacity. Here each line represents a different species. (c.ii) In each patch, individual survival probability decreases as the total community size increases. (d) Temperature changes stochastically over time. (d.i) Mean temperature decreases linearly with space. Over time, between (lower line) and (upper line), the temperature increases. (d.ii) Temperature variation over time depends on level of environmental stochasticity. The vertical dashed line designates when the model changes from the initialization phase (average temperature change ()) to the climate change and intervention phase (). Climate change only occurs after a relatively stable metacommunity has been assembled, after 100 time steps have passed with no extinctions. (e.i) Two types of fragmented environments compared: one with few large gaps and one with several narrow gap. (e.ii) Each of the four restoration management strategies (colored lines). Each involved increasing the integral of carrying capacity over space by an amount *E* more than the original carrying capacity (black lines). (f.i) Relocation occurs once the total population of a species falls below a threshold *η*. To avoid repetition while the species recovers, no relocations occur during a cool-down period following relocation *α*. (f.ii) A fraction *ρ* of the population is removed from its original distribution and moved to the closest new location where the average temperature 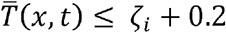 and the carrying capacity *K*(*x,y*) > 5 (only a fraction *μ* survive). Remaining individuals disperse naturally.

(Urban et al. 2012). To incorporate demographic stochasticity, the number of propagules produced by individuals in patch *x*, subpatch *y* is a Poisson random variable with mean equal to the reproductive output, 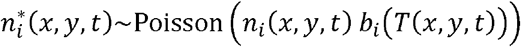 (Melbourne & Hastings 2008).

Next, each propagule disperses from its origin (Fig. 1b). Though reproduction occurs within the subpatch level, dispersal occurs at a larger spatial scale (between patches). Therefore, the model pools together all propagules in a patch prior to dispersal, such that the total number of propagules in patch *x* at time *t* is 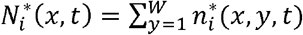. We adapt the Laplace dispersal kernel to a discrete-space analog, defining *λ*_*i*_, as the mean absolute distance (in patches) that individuals move from their origin and let kernel parameter 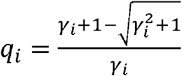. Thus, the probability of a propagule from patch *x* moving to patch *z* is

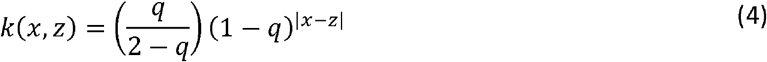

(Backus & Baskett 2021). All propagules disperse from patch *x* throughout all patches with a multinomial random vector. After arriving at patch *z*, propagules randomly distribute among the *W* subpatches of patch *z*. The resulting number of dispersed propagules in patch *z*, subpatch *y*, at time *t* is 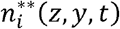.

Lastly, dispersed propagules compete for limited space and resources within each subpatch, given a location-dependent carrying capacity *K*(*x,y*) in each subpatch that remains constant over time (except when modified through management action) (Fig. 1c). The value of *K*(*x,y*) varies over space depending on the degree of habitat fragmentation. Density-dependent survival in this model is a variation on lottery competition (Sale 1978; Chesson & Warner 1981) with temperature dependence, with a higher chance of survival around a species thermal optimum *ζ*_*i*_ (Eq. 3). Altogether, each individual of species *i* has an equal probability of surviving,

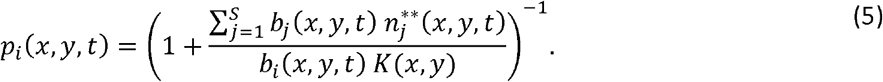

The total number of individuals that survive in patch *x*, subpatch *y*, after competition is a binomial random variable 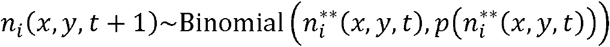 (Melbourne & Hastings 2008).

### Management interventions

We simulated six types of management strategies. Four of these strategies involved increasing the habitat quality in particular locations. To keep these strategies ecologically comparable, we increased the total carrying capacity by an amount defined as the “total area restored”, *E*. We let *K*_*u*_(*x,y*)be the unmanaged carrying capacity of patch *x*, subpatch *y*, and *K*_*m*_(*x,y*) be the carrying capacity after management. Then the total area restored is 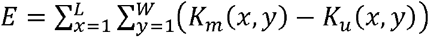.

With the “restore all” strategy, we increased the carrying capacity in all subpatches evenly by 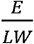. With the “corridor establishment” strategy, we increased the carrying capacity in all locations that were below a threshold carrying capacity and raised the minimum carrying capacity for all subpatches to that threshold. We numerically adjusted this threshold until the total area was *E*. With the “stepping-stone” strategy, we first identified all locations below a threshold. For each region with multiple patches below this threshold, we raised the carrying capacity for all subpatches in the middle 50% quantile of the gap but left the outer 25% quantiles at initial values. We adjusted this threshold until the total area restored was *E*. With the “reinforce” strategy, we increased the carrying capacity of all subpatches above a threshold, adjusting until the total area was *E*.

Following Backus & Baskett (2021), we simulated assisted migration by relocating species when the total metapopulation of a species falls below a threshold of *η* individuals (Fig. 1f). After the population of a species *i* fell below *η*, we relocated a fraction of the population *ϕ* to a location with a temperature approximately equivalent the species thermal optimum *ζ*_*i*_ in the future. To find this, we identified all locations with temperatures 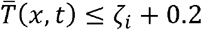. To avoid relocating a species into an area with low habitat quality, we only relocated the population into locations that fit the above specifications with *K*(*x,y*) *>* 5. We spread individuals between all subpatches within 5 patches (2 on either side of the target location). After relocating a population, we did not relocate that species again for *α* = 5 years to avoid relocating a population recovering from a previous relocation. Following optimal parameter values from Backus & Baskett (2021), we relocated *ϕ* of the total population and only *μ* survived relocation (Table 1). To limit assisted migration (to be somewhat comparable to habitat quality modification strategies), we only simulated relocations until we reached a maximum limit of *F*.

**Table 1:** Definitions of the symbols used in the model.

| Parameter | Symbol | Values | Units |
| --- | --- | --- | --- |
| Total species in pre-initialized community | $\Omega$ | 64 | species |
| Dispersal distance of species $i$ | $\gamma_i$ | Lognormal; mean=2.5, st. dev.=2.5 | patches |
| Thermal optimum of species $i$ | $\zeta_i$ | Uniform; 9.78 to 30.22 | °C |
| Thermal tolerance breadth of species $i$ | $\sigma_i$ | Lognormal; mean=5, st. dev.=5 | °C |
| Reproductive strength of species $i$ | $\rho_i$ | Derived from $\sigma_i$ | - |
| Skewness constant | $\lambda$ | -2.7 | - |
| Fraction of population relocated | $\phi$ | 0.55 | - |
| Assisted migration survival probability | $\mu$ | 0.8 | - |
| Low population threshold | $\eta$ | 50 or 75 | individuals |
| Cooldown time between relocations | $\alpha$ | 5 | years |
| Total patches | $L$ | 512 | patches |
| Subpatches per patch | $W$ | 8 | |
| Subpatch carrying capacity | $K(x, y)$ | Varies with space (average 8.25) | Individuals |
| St. dev. in local temperature heterogeneity | $H$ | Uniform; 0 to 2 | |
| St. dev. in interannual temporal stochasticity | $S$ | Uniform; 0 to 1 | |
| Mean annual temperature change | $\tau$ | 0.04 | °C/year |
| Annual temporal autocorrelation | $\kappa$ | 0.767 | - |
| Annual temporal standard deviation | $\psi$ | low=0.1639, high=0.6556 | °C |
| Total area restored | $E$ | $\frac{1}{8}LW$ to $LW$ (by $\frac{1}{8}LW$), $LW$ to $8LW$ (by $LW$) | individuals |
| Maximum relocations allowed | $F$ | 1 to 8 (by 1), 8 to 64 (by 8) | relocations |

### Numerical implementation

For our simulations, we used parameter values from Table 1. We used *L =* 512 patches and with *W =* 8 subpatches (a total of 2^12^ discrete locations). The initial mean temperature across the temperature gradient varied linearly from the poleward edge to the equatorward edge. Annual temporal autocorrelation was *k*, based on the measured combined global land-surface air and seasurface water temperature anomalies from 1880 to 1979 (GISTEMP Team 2019; Lenssen et al. 2019).

On average, the carrying capacity was a temperature-independent constant *K*(*x,y*) = 8.25 per subpatch so each patch could carry a total of 66 individuals at carrying capacity. In our simulations, we focus on two theoretical arrangements of high- and low-quality areas to represent different types of fragmentation: one with few wide gaps in habitat quality and one with several narrow (Fig. 1e). In each, the outer edges (*x* ≤ 64 and *x* ≥ 465) are at a constant intermediate carrying capacity *K*(*x,y*) *=* 8.25, while the center (65 ≤ *x* < 464) varies sinusoidally such that

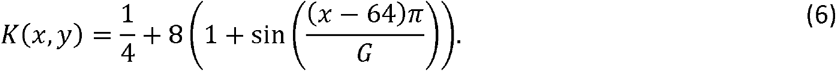

In environments with few wide gaps, *G =* 50, such that there are four full sine waves in the central region (spanning roughly 18.5°C of temperature change over space). In environments with several narrow gaps, *G =* 12.5, with 16 full sine waves in the central region.

In each set of simulations, we first generated the environment by randomly selecting the standard deviation of local heterogeneity *H* and environmental stochasticity *S* (Table 1). Next, we generated 64 species, selecting unique random values for each species’ thermal optima *ζ*_*i*_, thermal tolerance breadth *σ*_*i*_, and dispersal distance *γ*_*i*_. We numerically derived the reproductive strength *ρ*_*i*_, such that each species had the same overall reproductive potential *B =* 10 when integrating over temperature, emulating a jack-of-all-trades-master-of-none trade-off (Levins 1968). To generate the initial distribution and population size for all species in the community, we placed 4 individuals from all species in all subpatches and ran the model for 500 time steps with no change in average yearly temperature (*τ* = 0°C/year). At the end of this initialization phase, we used the final population sizes for each species in all subpatches as the initial conditions for climate change simulations.

Next, we simulated climate change on these initialized communities by adjusting the average yearly temperature change to *τ* = 0.04°C/year, roughly based on a “business-as-usual” projected scenario (Urban et al. 2012, IPCC 2021). This scenario provides the greatest number of extinctions with which to compare the relative efficacy of the different management strategies, where we expect that relative efficacy (the focus of our analysis) to remain consistent across different climate scenarios. For each community, we simulated the model for both 30 or 100 time steps after applying one of several management scenarios and degrees of management effort. In particular, starting at the beginning of the climate change (shift from *τ* = 0°C/year to *τ* = 0.04°C/year), we simulated “restore all”, “corridor establishment”, “stepping-stone”, and “reinforce” management strategies with total area restored values between 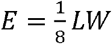 and *E = 8LW* (with 16 total variations; Table 1). Similarly, we simulated two threshold values for assisted migration (*η =* 50 or *η =* 75 individuals) with a maximum number of relocations between *F =* 1 and *F* = 64 (with 16 total variations; Table 1). For comparison, we also simulated community dynamics with no management effort (*E* = 0 and *F =* 0).

To evaluate how spatio-temporal heterogeneity affected management outcomes, we compared the number of extinctions prevented for corridor establishment and assisted migration (*η* = 75) under different levels of environmental stochasticity and local heterogeneity. To use comparable scenarios between these strategies, we chose values for *E* and *F* such that both strategies had a similar number of extinctions on average (*E* = 4*WL* for corridors and *F* = 8 for assisted migration). To evaluate which species benefited under the different management strategies, we found the extinction probability for each management action for species in each community that faced a variety of climate limitations: the species with the shortest average dispersal distance, the species with the narrowest thermal tolerance, the species with strongest competition in the poleward and equatorward direction (smallest difference in *ζ*_*i*_ values), and a random species for comparison.

## Results

Habitat corridors, stepping-stone reserves, and restoring all locations reduced the number of species that went extinct during climate change, and each of these strategies reduced extinctions further when restoring a larger total area (Fig. 2a,c). However, the reinforcing strategy had a negligible effect on extinctions. Both corridors and stepping-stones benefitted with relatively little area restored with diminishing returns with higher area restored, while restoring all locations reduced extinctions nearly linearly with increased area restored. On average, corridors reduced the number of extinctions more than all other restoration-based strategies with equivalent area restored. Stepping-stones reduced extinctions similarly to equivalent corridors with little area restored, but corridors were more effective than stepping-stones with higher area restored, especially in environments with fewer, larger gaps.

**Figure 2:**
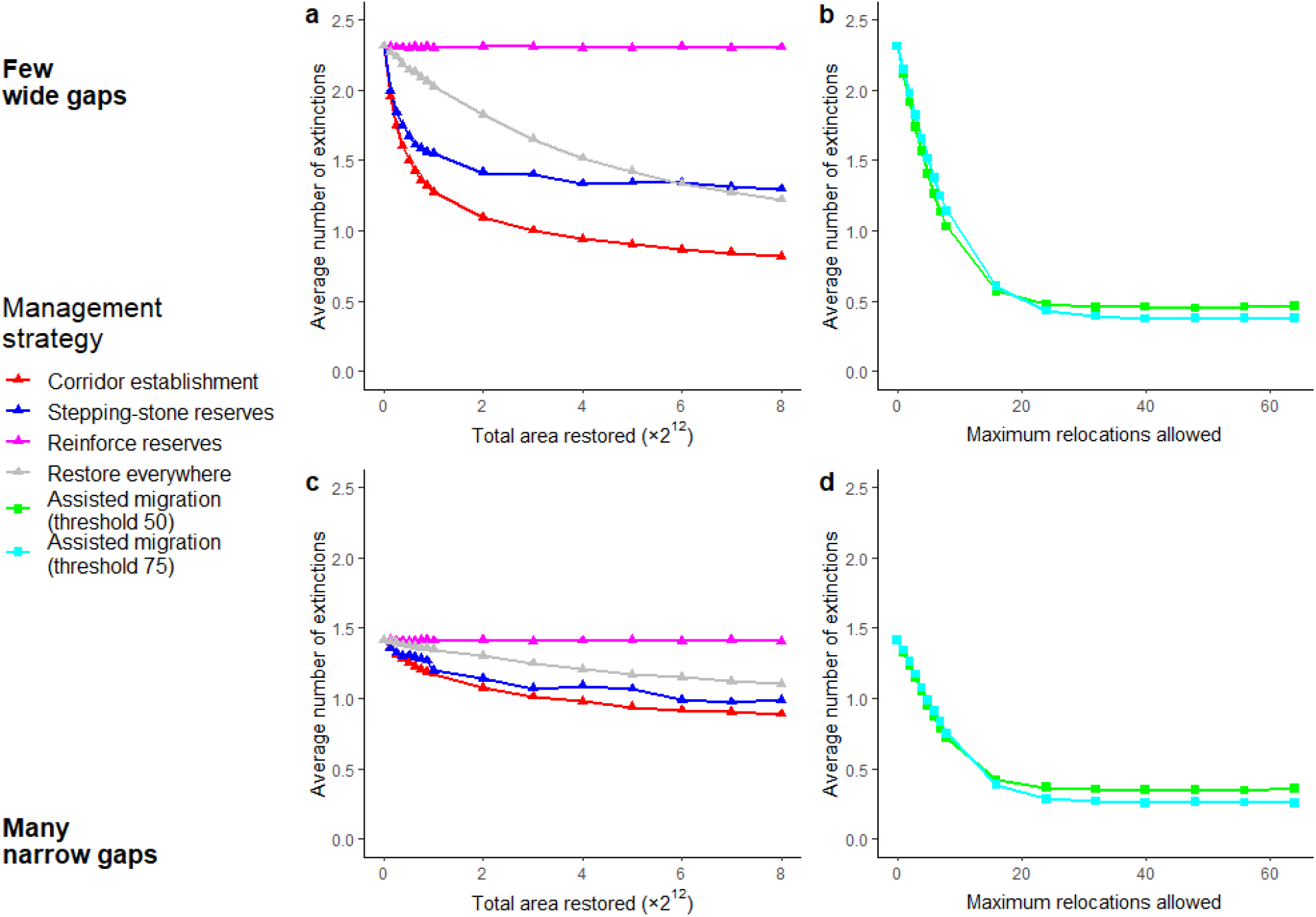
Average number of extinctions after 100 years (y-axis) in climate change simulations depending on management strategy (color/shape), amount of area restored for restoration-based modification (a,c) or maximum number of relocations allowed for assisted migration (b,d; x-axis), and environment structure (a-b: few wide gaps, c-d: many narrow gaps). Each point is the mean of 10000 simulations.

Assisted migration reduced extinctions on average, even with very few relocation events (Fig. 2b,d). However, increasing the maximum number of relocations above 16-24 did not reduce the average number of extinctions further. At this point, assisted migration prevented more extinctions on average than corridors at the highest area restored value we simulated. Both population thresholds for assisted migration that we simulated (*η* = 50 and *η =* 75) had similar extinction rates with equivalent relocation maximums.

Corridors were most effective at preventing extinctions in environments with low environmental stochasticity and moderate local heterogeneity (Fig. 3a,c), while assisted migration was most effective in environments with high heterogeneity and moderate stochasticity (Fig. 3b,d). Neither management strategy was effective at reducing the number of extinctions in environments with low heterogeneity and high stochasticity.

**Figure 3:**
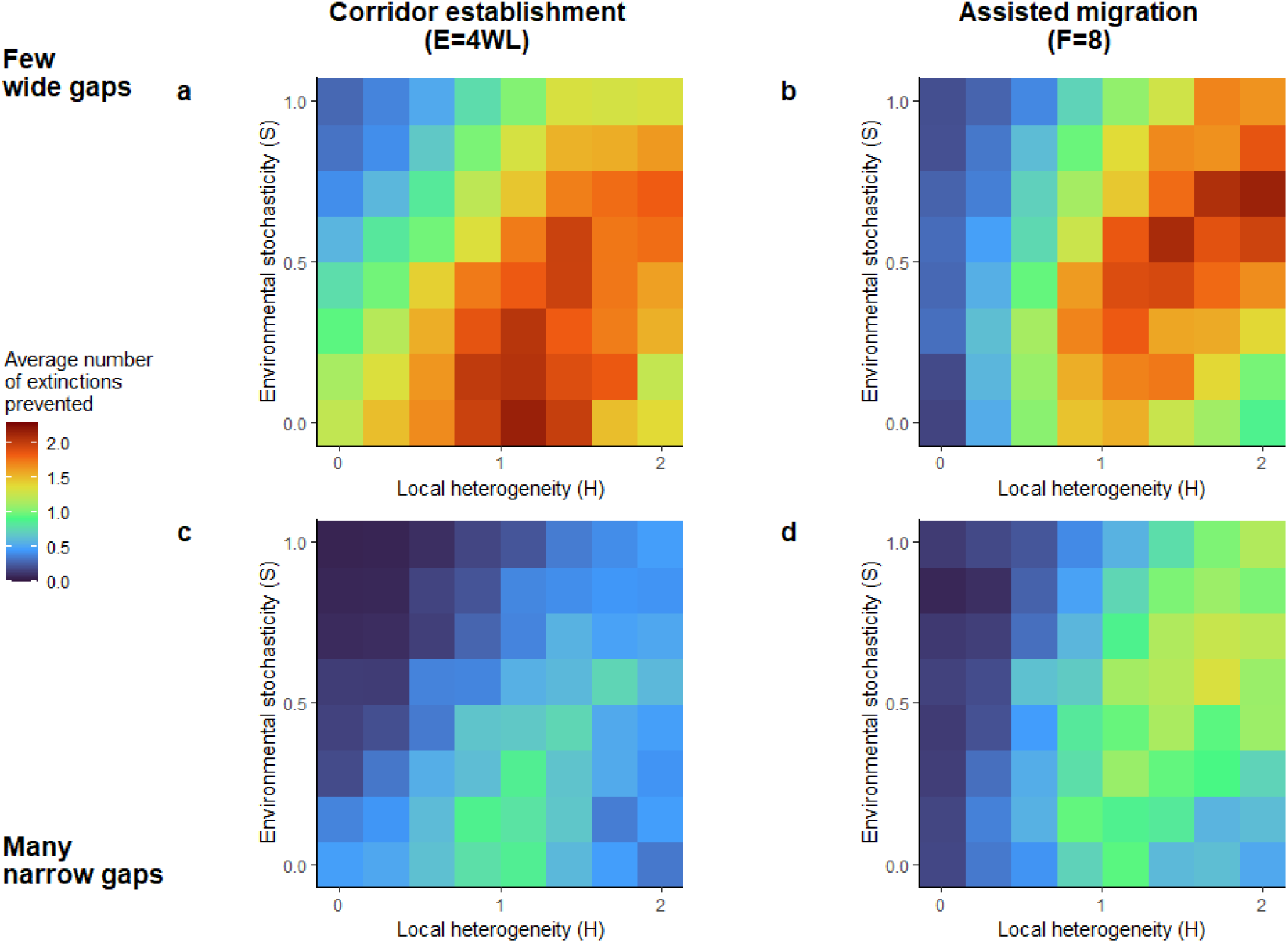
Average number of species prevented from extinction over 100 years (color) in climate change simulations as it depends on local heterogeneity *H* (standard deviation of temperatures per patch, x-axis) and environmental stochasticity *S* (standard deviation of interannual variation in temperature, y-axis). Each box represents the mean of between 135 and 189 simulations within an 8×8 quantiles of the range of all simulations. Panels (a) and (c) represent simulations with corridor establishment and panels (b) and (d) represent simulations with assisted migration. The total area restored in corridor establishment (*E* = 4*WL*) and the maximum number of relocations in assisted migration (*F* = 8) represent two cases where these two strategies prevent a similar number of extinctions on average for the “few wide gaps” environment, but not when comparing across equivalent levels of *H* and *S*. (a,b) represent simulations of environments with few wide gaps and (c,d) represent simulations of environments with several narrow gaps.

Randomly chosen species in simulated communities had a lower extinction probability under both corridor and assisted migration strategies, but the shortest dispersing species in a community disproportionately benefited more than random species (Fig. 4). Without management action, the shortest dispersing species had greater than 50% of going extinct throughout all variations of our simulations. Both management strategies reduced these extinction probabilities by more than 14% at similar effort levels (*E* = 4*WL* and *F* = 8). Reduction in extinction probability was greater for shortest dispersers than for random species in all scenarios. Other species likely to face extinction during climate change (narrowest thermal tolerance and the smallest difference in thermal optima with neighboring species on either pole- or equator-ward edges) were also less likely to face extinction with either management strategy, but only assisted migration reduced the extinction of these species disproportionately more than random species. Distinguishing the efficacy of assisted migration and corridors for different species and environmental conditions required longer-run (100 time step) simulations, as shorter-run (30-time step) simulations did not have enough extinctions to determine the impact of management interventions on extinction likelihood (2.0%-3.5% of species going extinct in 30 time steps *versus* 18.4%-31.6% of species going extinct in 100 time steps; Figs. S1-S2).

**Figure 4:**
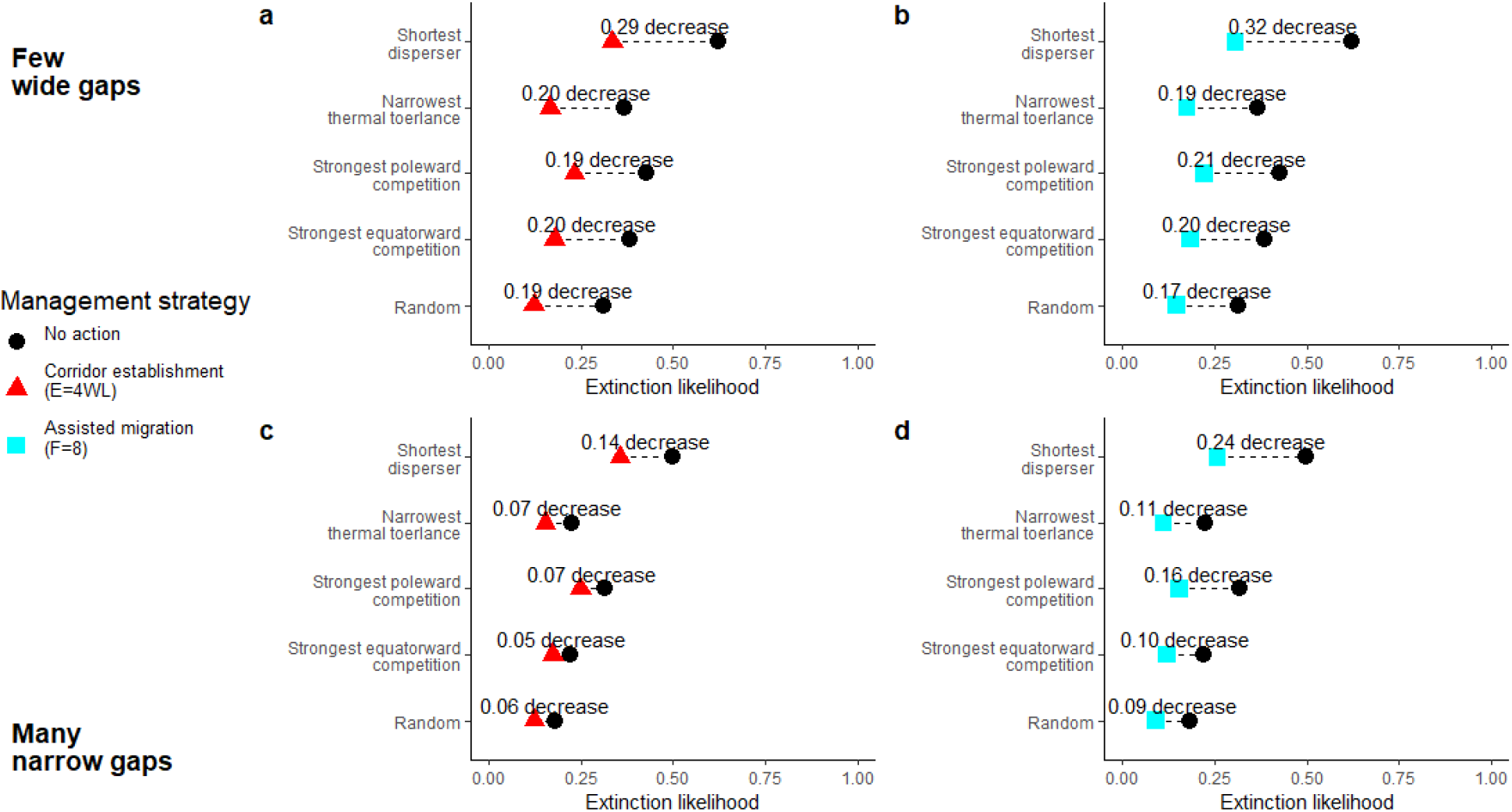
Likelihood that a species went extinct in 100 years (x-axis) in our climate change simulations depending on management strategy (color/shape, with corridors in panels a,c and assisted migration in panels b,d), which particular species it was in the community (y-axis) and environment type (a-b: few wide gaps, c-d: many narrow gaps). The particular species here are the species within the internal region of the environment (65 ≤ *x* < 464) with the shortest dispersal distance *λ*_*i*_, the species with the narrowest thermal tolerance *σ*_*i*_, the species in the community with extant neighboring species community closest to that species thermal optimum *ζ*_*i*_, and a randomly chosen species. Each point is the mean of 10000 simulations.

## Discussion

Most of the simulated management strategies reduced extinction probability under climate change in our simulated communities, and they reduced extinction rapidly with an initial investment in conservation effort. Without climate change, corridors, even when low quality, can facilitate species’ movement and long-term persistence in a metacommunity (Haddad & Tewksbury 2005; Williams et al. 2005). Adding to this, our model suggests that even relatively low-quality corridors between higher-quality areas could reduce extinction during climate change. Because restoring connectivity also increases total habitat area, the effects of increased connectivity and increased area are often confounded (Hodgson et al. 2009). Though many previous studies suggest that habitat reinforcement is often better at protecting species than connectivity restoration (Harrison & Bruna; Fahrig 2001; Falcy & Estades 2007), our results suggest that corridors are likely to be better at increasing the persistence of range shifting species in the presence of climate change than other methods of connectivity and protection that restore the same amount of area.

Similar to corridors, assisted migration reduced extinctions on average, even with relatively few relocation events in our model. Because many species in the simulated communities face little extinction risk from climate change, focusing relocation on a small number of vulnerable species was able to have a disproportionate effect on community-wide extinctions. If only a small number of species are at risk or conservation benefits can be realized by focusing on few species (Simberloff 1998; Enquist et al. 2020), but the few species at risk of extinction could require a high investment in management effort on their own. In practice, many conservation translocations are unsuccessful (Fischer & Lindenmayer 2000; Bubac et al. 2019), so managers might need to relocate a single species several times to increase the overall chance of establishment in the recipient location (Backus and Baskett 2021). Even after successfully establishing a new population, species with weak dispersal ability might continue to lag behind shifting climates and face extinction later. As climate change continues, these conservation-reliant species (*sensu* Scott et al. 2005) may depend on repeated direct management actions without increased connectivity (Lawler & Olden 2011).

Because we found relatively few extinctions in our nearer term simulations (30 time steps; Figs. S1-S2), the difference in the efficacy of management approaches was negligible, and we required long-run simulations (100 time step) to show the efficacy of corridors and assisted migration. This potential time lag to observable impact presents a challenge for monitoring to verify anticipated outcomes or adjusting management as needed in an adaptive management approach (Rist et al 2013). However, nearer-term impacts of management action might be evident in cases where optimal climates have already shifted away from species’ historical ranges, as has occurred for many species (Chen et al. 2011; Poloczanska et al. 2013), and our results suggest that near-term biodiversity conservation management can have long-term benefits for species persistence.

### Types of species benefitting from each management strategy

Adding to the extinction risks caused by fragmentation, many species are at risk of extinction from climate change because of a variety of biological limitations (Pearson 2006; Gilman et al. 2011; Urban et al. 2012; Urban 2015). We found that both corridors and assisted migration were effective at reducing the extinction of species with short dispersal in our model. These species benefited from increased connectivity regardless of the size of low-quality gaps. A previous simulation study showed that longer-dispersing competitors were likely to block shorter-dispersing species from tracking climate change in competitive communities with variable dispersal ability (Urban et al. 2012). Without connectivity, short-dispersing species that might disperse over patchy landscapes, but low population sizes, low propagule pressure, and strong competition means that these new populations are unlikely to establish (Lockwood et al. 2005; Plein et al. 2016). For corals, the species that are likely to have shorter average dispersal range, and likely to benefit from either corridor-like connectivity or assisted migration, are brooding species that release larvae directly from polyps rather than those that broadcast gametes into the water column (Ayre & Hughes 2000). Dispersal distance of trees is generally thought to be a function of seed size, tree height, and mode of dispersal (Bullock et al. 2016), where shorter trees that disperse seeds by wind or ballistics are more likely to have shorter dispersal than taller trees that dispersed seeds by birds.

In comparison, species narrow thermal tolerance and strong competition benefited more from assisted migration than restoration-based approaches. In corals, based on a trait-dependent clustering analysis of life history strategies, those with narrow thermal tolerance (i.e., outside of the “generalist” and “stress-tolerant” categories) and likely to experience strong competition (i.e., outside of the “competitive” category) fall into a category of “weedy” life histories, which are associated with small colony sizes and reproduction via brooding (where brooding increases reproductive success at low population sizes compared to mass spawning; Darling et al. 2012). Tree species with narrow geographic ranges may have narrow climate tolerance (though see Early & Sax 2014), whereas early successional species may face higher competition (Grime 1987).

Note that restoration and assisted migration are not dichotomous and can be integrated together in a larger management plan (Lawler & Olden 2011). Most tree species have low dispersal relative to climate change (Corlett & Westcott 2013), and most corals have narrow climate tolerance relative to climate change (Hughes et al. 2017), so these species could be threatened by climate change for multiple reasons. In these cases, increasing connectivity would benefit most species in the community and assisted migration would benefit those that disproportionately lag behind climate change.

### Environmental characteristics for different management strategies

In our simulations, the optimal management strategy depended on the characteristics of the environment. For example, species in environments with low stochasticity might especially benefit from corridor establishment over assisted migration. Because corridors are relatively small or low quality compared to the higher quality areas they connect, the population sizes in those corridors would be relatively small and susceptible to extinction (Lande 1993). Lower environmental stochasticity could allow a species to track climate change gradually, alongside several species competing to keep pace with climate change and move through the same limited area of a corridor. In coral reefs, one might identify regions of lower stochasticity through maps of past and projected degree heating weeks, a cumulative stress metric that predicts coral bleaching, which can then serve to inform the designation of reserve networks (Mumby et al 2011). In forests, one might preserve larger patches with smaller perimeter-to-area ratio, as edges between forest and fragments experience higher environmental stochasticity and frequency of rare weather events (Laurance 2004; Laurance et al. 2011).

In contrast, we found assisted migration to be particularly effective at reducing extinction in environments with moderate-to-high stochasticity. Because small populations are more likely to face extinction in environments with high environmental stochasticity (Lande 1993), both donor and recipient populations could face high extinction probability during assisted migration in stochastic environments. However, the benefits of moving a species near its optimal climate likely outweigh the risks of establishment failure on average, especially when planning multiple relocation events and relocating a fraction of a single population each time (Backus & Baskett 2021). Therefore, assisted migration might become an increasingly relevant management tool with increasing environmental variation and extreme events with climate change, such as marine heat waves in coral reefs (Fordyce et al 2019) and extreme droughts or fires affecting forests (Keeley & Syphard 2016; Williams et al. 2019). In our model, assisted migration was also more effective at reducing extinction in environments with higher local heterogeneity. Heterogeneous environments can act as climate refugia (Dobrowski 2011, Morelli et al. 2016), reducing the velocity of climate change or the negative effects of interannual variation. Because a highly heterogeneous recipient location is more likely to have a suitable microclimate for the relocated population to establish, relocating a population into a refugia-like environments could limit the risk of moving the population into the wrong place at the wrong time. For coral reefs, such local-scale heterogeneity and refugia might arise from fore-reef/back-reef structure, depth gradients, and physical structures that drive variability in local upwelling or tidal currents (Smith et al 2017). For forests, high local-scale heterogeneity is often found in areas with steep elevational gradients with similarly steep climate gradients (Morelli et al. 2016).

### Model assumptions

Even though a small amount of restoration or few relocations had large conservation benefits in our simulations, the actual economic and logistical costs of these strategies can be expensive. The total area restored metric does not fully reflect the economic costs of these approaches. To simplify comparison, we assumed that one unit of area restored (increasing the carrying capacity of the community by one individual) is equivalent for all species, regardless of how that area restored is distributed around the simulated environment. Realistically, conservation efforts and cost would vary across species and location (Naidoo et al. 2006; Magris et al. 2015). A corridor that spreads conservation spending across a wider range of low-quality areas would not be equivalent to a stepping-stone approach that uses the same spending in a smaller, condensed region. Also, considering inherent variation in land and water value or quality (Newburn et al. 2005), it would be difficult to improve the habitat quality of some locations, such as urban coastal waters, beyond a certain point. If the cost of protecting unbroken habitat corridors is prohibitive, “land sharing” approaches that allow conservation and human use to co-occur could enable connectivity (Green et al. 2005; Fischer et al. 2014).

Our model also simplifies some important ecological and evolutionary dynamics that might complicate comparisons between restoration and assisted migration-based approaches. In particular, our simple competition-based model does not include trophic interactions or disease dynamics. Incorporating these interactions could allow for relocated species to become invasive or spread disease, both of which are potential risks of assisted migration (Ricciardi & Simberloff 2009; Hewitt et al. 2011). We also ignore evolutionary dynamics in this model. Evolutionary dynamics could increase the effectiveness of connectivity-based approaches, as natural dispersal would favor increased gene flow of climate-tolerant genes as species naturally track climate change through corridors (Sgrò et al. 2011).

Lastly, we compared the extinction probability of species in our model, but other conservation goals might include maintaining ecosystem function or maintaining biomass for harvesting, among other goals. These alternative goals could favor different management strategies, as the benefits of each strategy are weighed by stakeholders depending on their willingness to engage in assisted migration with its high perceived risk or restoration-based approaches which could involve stakeholders giving up their land or harvesting rights. Further analysis of alternative management strategies to buffer against extinction from climate change and other conservation goals would benefit from a structure-decision making approach that considers the full array of risks, benefits, and uncertainties related to the array of potential stakeholder goals.

## Acknowledgments

We thank L. Bay, C. Clements, R. Gates, S. Harrison, C. Logan, M. McClure, C. Muhlfeld, S. Sawyer, M. Schwartz, R. Waples, and A. Whipple for their thoughtful conversations at the managed relocation workshop at UC Davis that informed this manuscript.

## Data accessibility

Simulation code, simulation results, and code to reproduce the plots in this paper are available at https://github.com/gabackus/comparingManagementStrategies.

