## supplementary figures for "Comparing management strategies for conserving communities of climate-threatened species with a stochastic metacommunity model"

Supplementary material – Management comparisons after 30 years of simulation

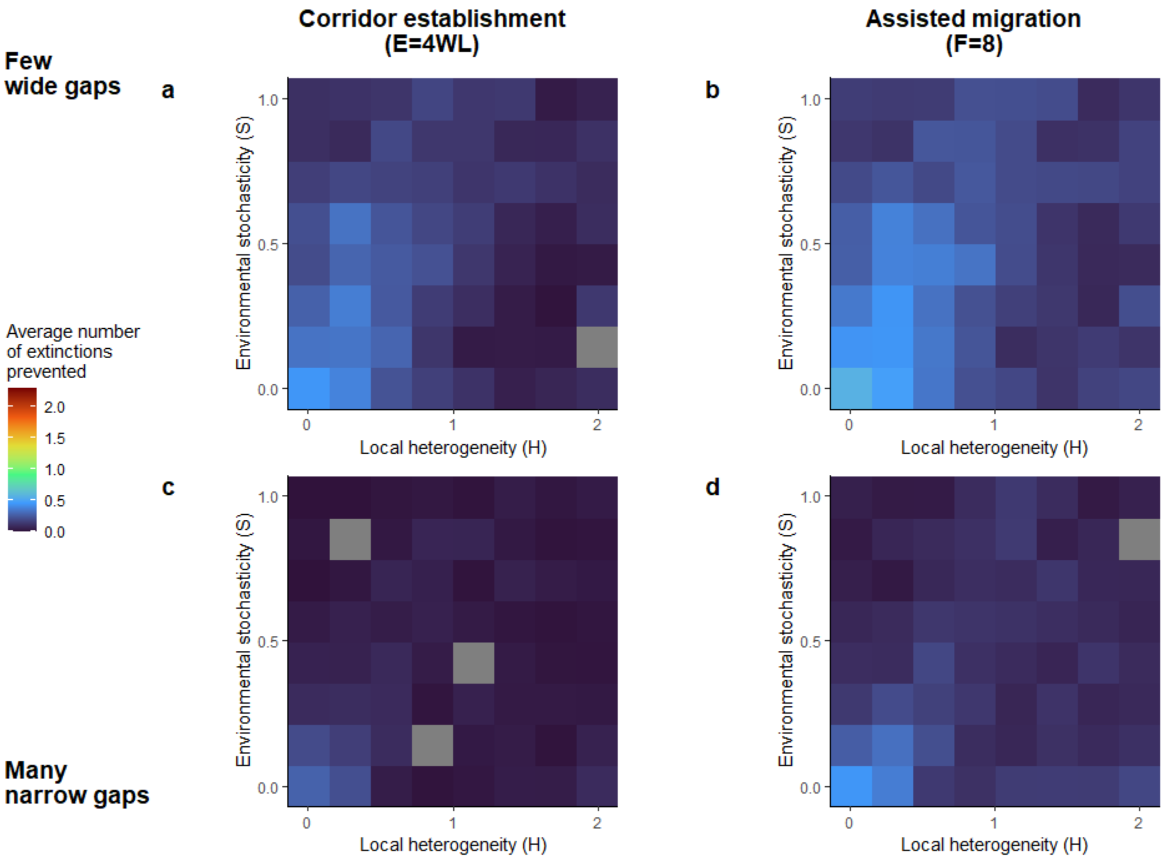


Figure S1: Average number of species prevented from extinction over 30 years (color) in climate change simulations as it depends on local heterogeneity $H$ (standard deviation of temperatures per patch, x-axis) and environmental stochasticity $S$ (standard deviation of interannual variation in temperature, y-axis). Each box represents the mean of between 135 and 189 simulations within an 8x8 quantiles of the range of all simulations. Panels (a) and (c) represent simulations with corridor establishment and panels (b) and (d) represent simulations with assisted migration. The total area restored in corridor establishment ($E = 4WL$) and the maximum number of relocations in assisted migration ($F = 8$) represent two cases where these two strategies prevent a similar number of extinctions on average for the "few wide gaps" environment, but not when comparing across equivalent levels of $H$ and $S$. (a,b) represent simulations of environments with few wide gaps and (c,d) represent simulations of environments with several narrow gaps.

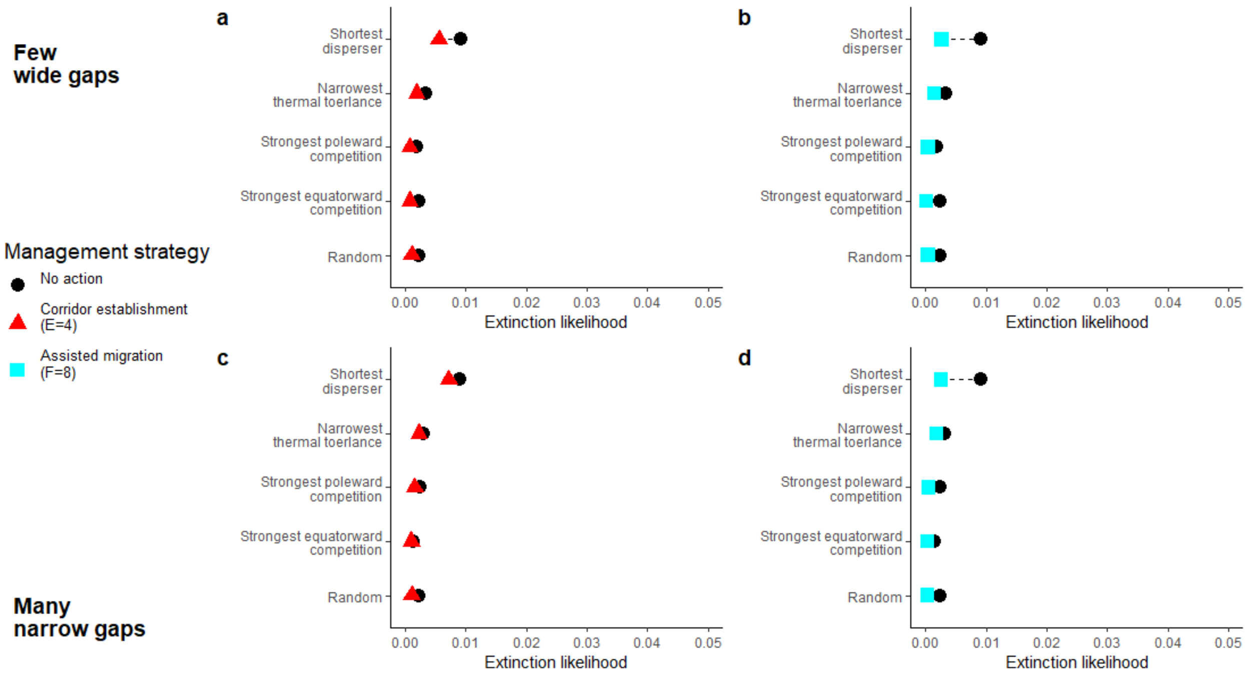


**Figure S2**: Likelihood that a species went extinct in 30 years (x-axis) in our climate change simulations depending on management strategy (color/shape, with corridors in panels a,c and assisted migration in panels b,d), which particular species it was in the community (y-axis) and environment type (a-b: few wide gaps, c-d: many narrow gaps). Note that the x-axis shows a much smaller range than a similar Figure 4 in the main text. The particular species here are the species within the internal region of the environment ($65 \leq x < 464$) with the shortest dispersal distance $\gamma_i$, the species with the narrowest thermal tolerance $\sigma_i$, the species in the community with extant neighboring species community closest to that species thermal optimum $\zeta_i$, and a randomly chosen species. Each point is the mean of 10000 simulations.
